# Prevalence of hemic neoplasia in cockles and co-habiting bivalve species in the western coast of France and the southern coast of Portugal

**DOI:** 10.1101/2025.07.30.667629

**Authors:** Alicia L. Bruzos, Angèle Moulin, Géo Bujard, Anthony Sturbois, Alejandro Viña-Feás, Ana Margarida Amaral, Camille Détrée

## Abstract

Hemic neoplasia is a cancer affecting several bivalve species and has become transmissible in some lineages. This study assesses its prevalence in common cockles (*Cerastoderma edule*) along the western coast of France and the southern coast of Portugal, regions with important bivalve fisheries. Cockles and cohabiting bivalve species (*Ruditapes* spp. and *Scrobicularia plana*) were sampled across multiple locations, and hemic neoplasia was diagnosed through cytological analysis. In the Algarve, a decline in prevalence was observed between 2017 and 2024, while in Normandy, the disease was detected for the first time. No cases of hemic neoplasia were found in the cohabiting clam species, suggesting that cockle cancer transmission may be species restricted. Morphometric analyses revealed no significant differences in size or weight between affected and non-affected cockles, indicating that external traits are not reliable indicators of infection. These findings expand knowledge on the geographic distribution of hemic neoplasia and highlight the need for continued monitoring to assess potential changes in its prevalence and impact on bivalve populations.

## 1. INTRODUCTION

Cancer is a group of diseases characterized by uncontrolled cell growth and proliferation, which can lead to tissue invasion and, in some cases, metastasis to other parts of the body (Stratton, Campbell and Futreal, 2009). Cancers have been observed across the tree of life (Aktipis *et al*., 2015) from most vertebrates to echinoderms (starfish, sea urchins…), cnidaria (jellyfish, corals, anemones…) or bivalves (cockles, clams, mussels…). Unlike many diseases caused by infectious agents such as bacteria or viruses, cancer is generally considered a non-contagious disease, originating from somatic mutations in an individual’s cells (Stratton, Campbell and Futreal, 2009). However, there are rare exceptions where cancer can be directly transmitted between individuals. These transmissible cancers, though uncommon in vertebrates, are well-documented in certain marine bivalve species (Ní Leathlobhair and Lenski, 2022).

Transmissible or contagious cancers are clonally derived cell lines that can spread from one individual to another via the transfer of living cancer cells, thus surviving beyond the death of the individuals that spawned them and effectively behaving like infectious diseases (Murchison *et al*., 2010). While certain cancers are initiated by infectious agents such as viruses (e.g., HPV) or bacteria, these are not transmissible as clonal cancer cell lines; in contrast, transmissible cancers involve the direct transfer of living neoplastic cells between individuals. In mammals, such cancers are limited to a few known examples, such as the canine transmissible venereal tumour in dogs and the Tasmanian devil facial tumour disease (Ní Leathlobhair and Lenski, 2022). In contrast, several species of marine bivalves have been found to be affected by naturally occurring transmissible cancers (Metzger *et al*., 2015). These diseases are typically hemic neoplasias, affecting haemocytes and have been found across the bivalve tree in distantly related lineages such as mussels, clams and common cockles (Metzger *et al*., 2015, 2016; Yonemitsu *et al*., 2019, 2025; Hammel *et al*., 2021; Garcia-Souto *et al*., 2022; Michnowska *et al*., 2022). Bivalve transmissible neoplasia (BTN) is associated with disruption of physiological functions and can lead to population declines when prevalence is high (Carballal *et al*., 2015). The disease is of particular concern in aquaculture regions as epizootiological outbreaks have been reported in association with mass mortalities in various bivalve species (Da Silva *et al*., 2005).

Among the bivalve species affected by transmissible hemic neoplasia, the common cockle (*Cerastoderma edule*) has emerged as one model for studying the disease. Cockles are widely distributed along European Atlantic coast, from Rusia to Senegal, and play an essential role in intertidal ecosystems (Maia, Barroso and Gaspar, 2021). This species lives buried under the surface in clean sand, muddy sand, or muddy gravel bottoms, and it is commonly found in intertidal flats and shallow subtidal areas of estuaries, coastal lagoons and sheltered coastline bays. Their filter-feeding activity contributes to the cycle of nutrients and sediment stability (Dabouineau et *al*., 2015). Additionally, cockles represent a valuable economic resource in commercial fisheries and aquaculture (Carss *et al*., 2020). Several studies have identified hemic neoplasia outbreaks in cockles across multiple European regions, with varying prevalence levels (Barber, 2004; Romalde *et al*., 2007; Le Grand *et al*., 2010; Ruiz *et al*., 2013; Montaudouin *et al*., 2021). Notably, in 2017, the highest reported occurrence was found in the Algarve, southern Portugal, among 36 locations screened along the European Atlantic coast (Bruzos *et al*., 2023).

A critical aspect of transmissible cancer in bivalves is its potential for cross-species transmission. Unlike most cancers, which originate within a single host and do not spread between individuals, some transmissible cancers in bivalves have been found in multiple species (Metzger *et al*., 2016; Yonemitsu *et al*., 2019; Hammel *et al*., 2021; Garcia-Souto *et al*., 2022). This phenomenon raises important ecological and epidemiological questions regarding the mechanisms underlying interspecies transmission and the potential for marine environments to facilitate such events. Cohabitation of different bivalve species within the same habitat provides opportunities for cell transfer through water, potentially increasing the risk of disease spread (Giersch *et al*., 2022; Burioli *et al*., 2021).

Previous research has documented three cases of interspecies transmission of BTN. The BTN observed in *Polititapes aureus* originated in a related species, *Venerupis corrugata*, despite only sporadic cases of BTN being found in *V. corrugata*, suggesting a potential resistance in which the cancer has persisted by engrafting into a new host species (Metzger *et al*., 2016). Similarly, the BTN observed in *Venus verrucosa* originated in *Chamelea gallina* and no cases have been reported in the later (Garcia-Souto *et al*., 2022). Lastly, one of the two BTN lineages identified in *Mytilus trossulus* has successfully spread to three additional mussel species and hybrids, highlighting the complexity of transmissible cancer propagation in bivalves (Yonemitsu *et al*., 2019; Hammel *et al*., 2021; Skazina *et al*., 2021).

To date, BTN reports have been made in coastal locations from four continents and all oceans. These global inter-and intraspecific contagions illustrate the urgency of identifying and characterizing these contagious cancers in bivalve species in order to prevent their spread and manage the aquaculture production (Hammel *et al*., 2024).

Portugal’s bivalve fisheries, particularly in the Algarve, represent one of the most economically significant shellfish industries in Europe (Carss *et al*., 2020). The Algarve is a major hub for bivalve aquaculture, with extensive production of clams (*Ruditapes decussatus* and *R. philippinarum*), cockles (*C. edule*), and other species within the Ria Formosa lagoon and close coastal waters (Brito *et al*., 2012). This region’s warm climate and productive estuarine habitats support high shellfish yields, making it a cornerstone of local fisheries and export markets (Carss *et al*., 2020). However, the occurrence of transmissible hemic neoplasia in cockles poses a threat to the stability of these populations (Bruzos *et al*., 2023). Understanding the disease’s prevalence and potential for spread is essential for ensuring sustainable fisheries management and mitigating economic losses in the region.

Similarly, in France, the Atlantic coastline is home to significant bivalve populations, particularly in regions such as Brittany and Normandy (Dabouinneau et *al*., 2015). These areas have long-standing traditions of shellfish harvesting, with oysters, mussels, and clams playing an important role in local economies (Carss *et al*., 2020). The presence of hemic neoplasia in French cockle populations has been relatively understudied, though cases have been documented in Brittany (Poder and Auffret, 1986; Le Grand *et al*., 2010; Bruzos *et al*., 2023). This study aims to provide the first systematic assessment of hemic neoplasia in cockles along the Normandy coast, expanding our knowledge of the disease’s distribution and potential ecological impact.

Beyond disease prevalence and epizootiological outbreaks, this study aims to investigate the relationship between hemic neoplasia and morphometric traits in cockles. Parasitic infections in bivalves have been linked to changes in growth rates, body condition, and overall fitness (Zannella *et al*., 2017). Morphometric analysis can provide valuable insights into whether affected individuals exhibit distinct physical characteristics compared to their healthy counterparts. For that, we integrate morphometric data with hemic neoplasia assessments to explore potential correlations between disease status and shell size and weight in cockles from both the Algarve and Normandy.

By investigating the prevalence and morphometrics of hemic neoplasia in cockles and cohabiting bivalve species across the southern coast of Portugal and the western coast of France, we aim to improve our spatio-temporal understanding of BTN’s epidemiology.

## 2. METHODOLOGY

### 2.1. Target species selection and animal collection

The primary target species for this study was *Cerastoderma edule* (common cockle, Figure 1A). We also selected the cohabitant bivalves (Figure 1B-C) *Scrobicularia plana* and *Ruditapes* spp. (including *R. decussatus* and *R. philippinarum*, which cannot be visually distinguished). These species were chosen based on three main factors: (1) they share ecological habitats with *C. edule*, (2) they are closely related evolutionarily, and (3) they have significant economic value in aquaculture and can be easily collected from the same environments (Carss *et al*., 2020). A total of 930 bivalve individuals were collected, including 460 *Cerastoderma edule* (267 from Brittany/Normandy and 193 from the Algarve), 295 *Ruditapes* spp. (14 from Normandy and 281 from the Algarve), and 175 *Scrobicularia plana* (19 from Brittany and 156 from the Algarve).

**Figure 1.**
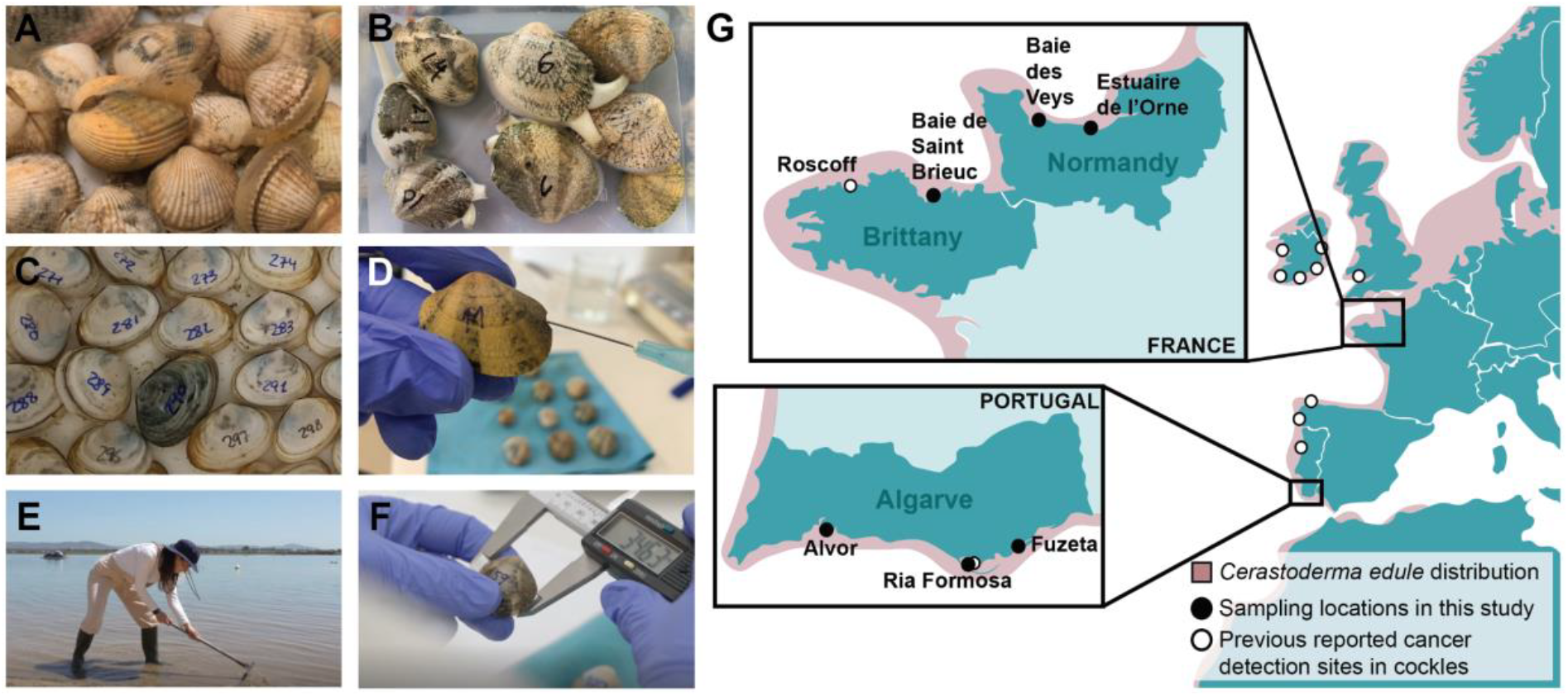
Bivalve species, sampling locations, and methods. **(A)** Common cockle (*Cerastoderma edule*), **(B)** Palourde clam (*Ruditapes* spp.), **(C)** *Scrobicularia plana*. **(D)** Hemolymph extraction from the adductor muscle using a syringe with a needle. **(E)** Manual collection of cockles from the sand using a rake (photograph depicts A. L. Bruzos, author of this manuscript). **(F)** Measurement of bivalve size with a digital caliper. **(G)** Map of the Brittany / Normandy and Algarve regions showing sampling locations from this study and previously reported cancer detection sites in cockles.

### 2.2. Field sampling and sample maintenance

Sampling was conducted at 6 locations of two regions (Figure 1G): Algarve (South of Portugal) and Northern Brittany/Normandy (West of France). The Algarve was chosen due to its high prevalence of hemic neoplasia in cockles reported in 2017 (Bruzos *et al*., 2023), and the economic importance of this aquaculture region within Europe (Carss *et al*., 2020). Normandy was chosen due to its proximity to Brittany, where cockles were previously identified with hemic neoplasia, though Normandy itself had no prior reports of such cases. Specific locations were selected based on previous studies (Quéro *et. al*, 1998) with the support of local shell-fishers and technicians.

Sampling was carried out in three time periods considering weather and tidal conditions: summer 2023 for Brittany, spring 2024 for Normandy, and summer 2024 for the Algarve. Cockles of different sizes were collected from natural intertidal sand beds using manual sampling methods (Figure 1E) to ensure representative sample sizes for prevalence assessment. The number of collected cockles varied slightly due to natural differences in cockle populations, but a minimum of 30 individuals per site was targeted for each sampling location. For cohabiting species, specimens of *Ruditapes* spp. and *S. plana* were collected from the same sand beds when present. Consequently, the number of individuals varied depending on their local availability.

Samples were maintained alive in closed-circuit seawater tanks for 48 hours prior to diagnosis, measurements and additional procedures. Animals from different sampling locations were never mixed within the same tank, and tanks were thoroughly cleaned with bleach between sample arrivals to prevent cross-contamination. Samples collected in Normandy and Brittany were transported to the Centre de Recherches en Environnement Côtier (CREC) at Université de Caen Normandie in Luc-sur-Mer, France, while those from the Algarve were kept at the Ramalhete Marine Station, Centro de Ciências do Mar (CCMAR) in Faro, Portugal.

### 2.3. Morphometric measurements and statistical analyses

Bivalve specimens were counted, measured with a digital calliper (precision of 0.01 mm; Figure 1F), and weighed using a top-loading digital balance (precision of 0.01 g). Morphometric analysis included two primary shell dimensions: shell length (SL, the maximum anterior-posterior distance) and shell height (SH, the maximum dorsal-ventral distance across the middle axis of the shell). Total weight (WE) was determined after briefly blotting specimens to remove surface fluids and draining excess water from the mantle cavity. Dry weight was not measured to avoid harming the bivalves required for diagnostic protocols. Damaged specimens, mainly those with anthropogenic damage from fishing and handling, were either discarded or only partially analysed, depending on the intactness of their morphometric parameters.

To further characterize the morphology, two morphometric indices were calculated: the elongation index (SH/SL) and the density index (WE/SL), using ratios between shell dimensions and weight following the equations of (Caill-Milly *et al*., 2012, 2014).

Statistical analyses were conducted using R (version 4.4.2). Morphometric differences between cancerous and non-cancerous cockles were assessed through Student’s t-tests or Mann-Whitney U tests, depending on normality (Shapiro-Wilk test). Pearson’s correlation coefficients were computed to examine the relationships among weight, length, and height, with separate analyses for cancerous and non-cancerous individuals. Violin plots and pairwise scatter plots were generated using the *ggplot2* (Wickham, 2016) and *ggally* (Schloerke, 2025) packages to visualize morphometric distributions and correlations.

### 2.4. Cytological diagnosis

Hemic neoplasia was diagnosed through examination of haemolymph cell monolayers. Although cytological examination allows for reliable detection of hemic neoplasia, it does not enable confirmation of lineage identity; therefore, transmissibility cannot be inferred without complementary molecular analysis. Haemolymph from each bivalve was extracted following a non-invasive technique based on Vandepas *et al*. (2023). For *Cerastoderma edule* and *Ruditapes* spp., a small hole was made in the shell using a fine file to facilitate syringe access (Figure 1D), while *Scrobicularia plana* specimens were accessible without shell modification.

Haemolymph was withdrawn from the adductor muscle of each bivalve sample using a 23-gauge needle attached to a 5 ml syringe. To prevent haemocyte aggregation, 50 µL of haemolymph were mixed with 150 µL of cold modified Alsever’s anti-aggregate solution (Bachère, Chagot and Grizel, 1988). This mixture was gently homogenized to avoid cell rupture, then pipetted onto a microscope slide and air-dryed for 8 to 15 minutes. Slides were examined under a microscope using a Leica DM2000 LED light microscope in France and with a ZEISS Primo Star light microscope in Portugal. Positive cases were subsequently fixed and stained with the kit Hemacolor (Merck) and reviewed.

Prevalence rate was calculated as the percentage of positive cases out of the total number of individuals examined per location, year and species. Disease severity in cockles was categorized based on a scale by manually counting 100 cells per sample: non-affected (N0) when no cancer cells were observed, early-stage cancer (N1) when cancer cells were under 15%, medium-stage cancer (N2) when 15-75%, and severe-stage cancer (N3) when above 75% (Díaz et al., 2010). Positive samples were preserved in ethanol for future molecular analysis.

### 2.5. Ethical considerations and permissions

All sampling activities complied with local and European regulations on marine research. Permits for collecting biological materials were obtained from authorities in Portugal and France, including specific permissions coordinated by the CCMAR research manager for sampling within Ria Formosa Natural Park. Animal procedures were planned according to European Directive 2010/63/EU and Portuguese and French legislation. To minimize impact, the number of collected bivalves was kept low, targeting only 1-5 diseased specimens per site, and diagnostic protocols were optimized to limit harm, requiring just 50 µL of haemolymph per individual.

## 3. RESULTS

### 3.1. Temporal evolution of the prevalence of Hemic Neoplasia in the Algarve

In 2017, cockles from the Algarve showed the highest prevalence of hemic neoplasia among 36 locations studied across 11 countries (Bruzos *et al*., 2023). To assess the evolution of the prevalence of the disease in the region, we analysed data over three years: 2017, 2020 and 2024 (Figure 2A-B). In 2017, cockles from Ria Formosa presented a prevalence rate of 23.1% (72 of 312 individuals). By 2020, prevalence increased to 31.4% (80 of 255 individuals), indicating a potential rise in cancer incidence over the three-year period. However, in 2024, the trend reversed, with prevalence declining to 9.3% (7 of 75 individuals). A Fisher’s exact test comparing prevalence across the three years indicated that the differences were statistically significant (2017 vs 2020: *p* = 0.029; 2017 vs 2024: *p* = 0.007; 2020 vs 2024: *p* = 0.00008). The number of cockles analysed decreased over time due to increasing difficulty in locating individuals in Ria Formosa. This decline in abundance is consistent with observations from local shell-fishers, who have reported a reduction in cockle populations over the past decade (personal communication). Given the difficulty in gathering 100 cockles from the sampling site, we added two nearby locations for the 2024 study (Figure 2C, Table 1): Fuzeta, located 11.8 nautical miles east of Ria Formosa, showing a prevalence of 8.8% (3 of 34 individuals), and Alvor, 31.4 nautical miles west, where no case was detected (0 of 84 individuals). Regarding the stages of cancer, a higher number of samples were detected at stage N2 or N3 in all sampling locations (Figure 2D) compared to previous reports (Bruzos *et al*., 2023).

**Table 1.**
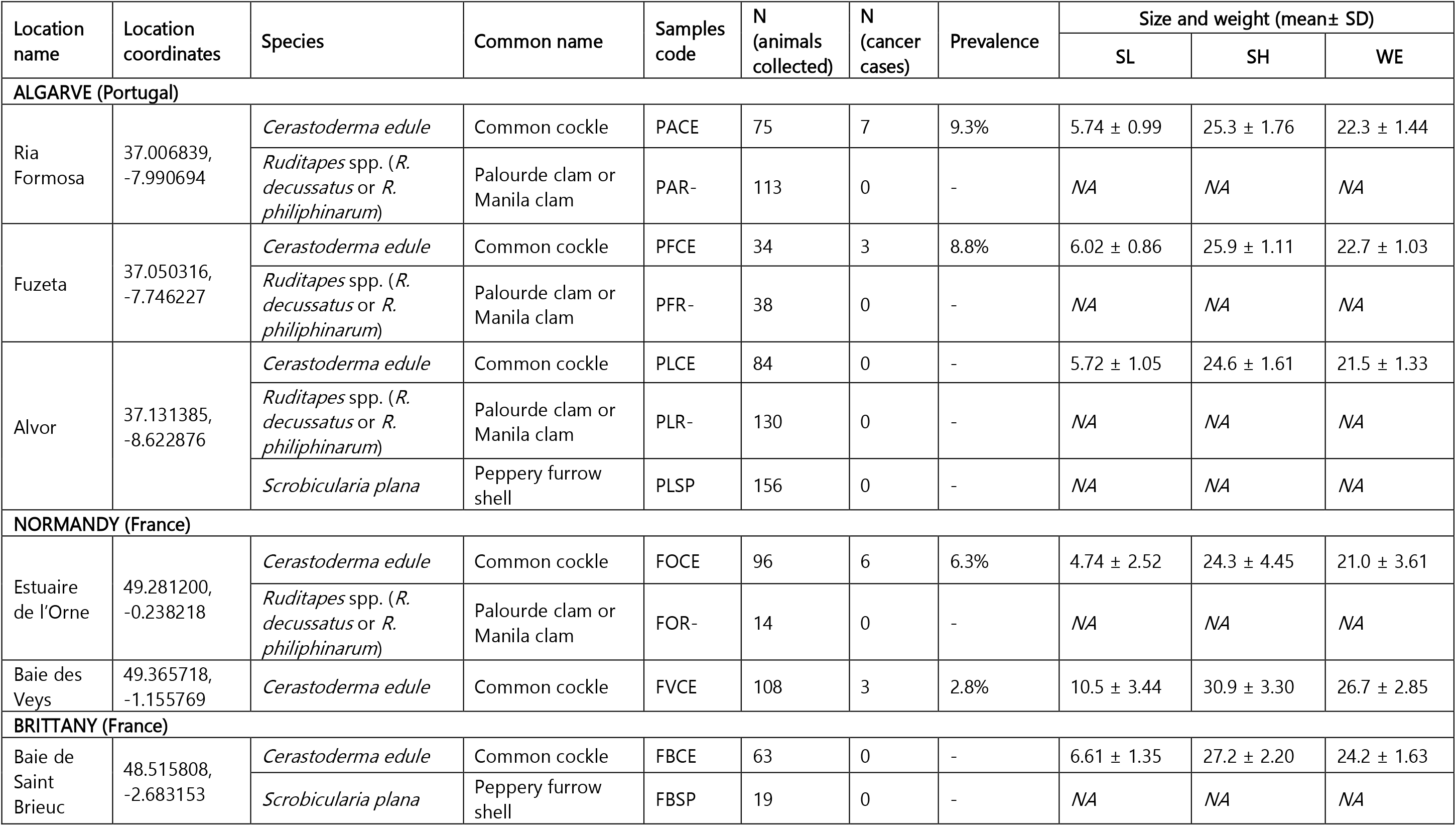
Descriptive statistics, sampling locations, morphometric measurements, and cancer diagnoses for three common bivalve species from the Algarve coast (southern Portugal) and the Normandy and Brittany coast (western France).

**Figure 2.**
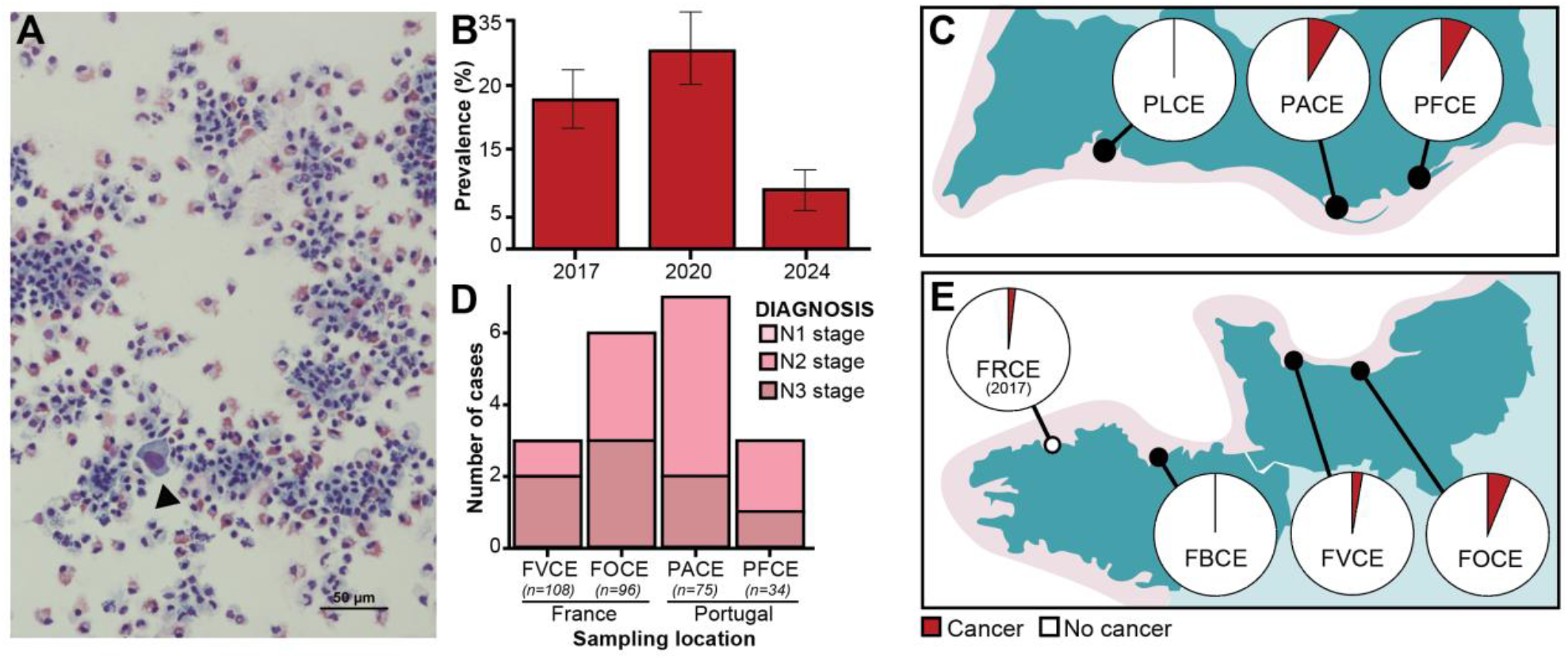
Variations of hemic neoplasia prevalence in cockles. **(A)** Cytological preparation of haemolymph from a cockle of the Algarve, arrowhead points a cancer cell, bar 50 μm. **(B)** Temporal evolution of cancer prevalence in Ria Formosa (Algarve, Portugal) over three years: 2017, 2020, and 2024. **(C)** Piecharts of cancer prevalence of across the Algarve (Portugal) in 2024. **(D)** Distribution of cancer stages, represented as stacked bar plots; n indicates the number of collected animals. **(E)** Piecharts of cancer prevalence across sampling sites in Normandy and Brittany (France). For comparison, data from Roscoff (FRCE from Bruzos et al., 2023) is included.

### 3.2. Hemic Neoplasia detection in common cockles from Normandy

This study is the first to document hemic neoplasia in cockles along the Normandy coast of France. Previously, cases had only been reported in the neighbouring Brittany region, particularly in Roscoff, where Bruzos et al. (2023) reported a prevalence of 2.1% (5 of 240) in 2017 and 1.4% (2 of 144) in a second sampling that year.

In Normandy, hemic neoplasia was detected at two locations (Figure 2E). In the Estuaire de l’Orne, prevalence reached 6.3% (6 of 96 individuals), while in Baie des Veys, it was 2.8% (3 of 108 individuals). Interestingly, in the nearby Brittany sampling site of Baie de Saint Brieuc, which lies midway along the coastline between Roscoff and the Normandy studied sites, no cancer cases were detected (0 of 63 individuals). No sample was detected at the initial stage in any of the sampling locations in Normandy. This new detection in Normandy extends the known range of hemic neoplasia in French cockle populations, highlighting its spread along the northern coast of France.

### 3.3. Assessment of cross-species transmission to co-habiting clam species

As cross-species transmission of hemic neoplasia has already been detected in several species, to investigate the potential risk for cohabiting bivalve species within the areas where we detected hemic neoplasia, we sampled two bivalve species cohabiting with cockles in both the Algarve and Normandy regions. A total of 295 *Ruditapes* spp. (including *R. decussatus* and *R. philippinarum*) and 175 *S. plana* individuals were examined and no evidence of hemic neoplasia in any individuals from either species was found.

### 3.4. Morphometric analysis

Morphometric analysis was performed on all bivalves collected to explore any potential association between cancer presence and alterations in size or weight. The morphometric analysis did not reveal significant differences between cancerous and non-cancerous cockles in terms of weight, length, or height (Table 2). As shown in the violin plots (Figure 3A), the distributions of these measurements were largely overlapping, with no clear separation between infected and non-infected individuals. This pattern was consistent across both Portuguese and French populations, indicating that cancer status did not have a strong effect on overall shell morphology or body weight.

**Table 2.** Comparison of morphological measurements between cancerous and non-cancerous individuals. Values are presented as mean ± standard deviation (SD).

| Measurement | Population | Cancer<br>(mean± SD) | No cancer<br>(mean± SD) | Wilcoxon<br>p value | Interpretation |
| --- | --- | --- | --- | --- | --- |
| Weight (WE) | All | 6.62 ± 2.64 | 6.80 ± 3.08 | 0.849 | No significative |
| | Portugal | 5.93 ± 1.22 | 5.77 ± 0.984 | 0.803 | No significative |
| | France | 7.38 ± 3.58 | 7.54 ± 3.77 | 0.859 | No significative |
| Length (SL) | All | 26.6 ± 3.09 | 26.6 ± 3.90 | 0.876 | No significative |
| | Portugal | 25.5 ± 1.52 | 27.7 ± 4.61 | 0.532 | No significative |
| | France | 27.7 ± 4.01 | 25.1 ± 1.67 | 0.920 | No significative |
| Height (SH) | All | 23.1 ± 2.59 | 23.2 ± 3.25 | 0.942 | No significative |
| | Portugal | 22.5 ± 1.49 | 24.1 ± 3.88 | 0.538 | No significative |
| | France | 23.8 ± 3.39 | 22.0 ± 1.40 | 0.797 | No significative |
| Elongation<br>index (SH/SL) | All | 0.87 ± 0.03 | 0.88 ± 0.04 | 0.518 | No significative |
| | Portugal | 0.88 ± 0.03 | 0.88 ± 0.05 | 0.836 | No significative |
| | France | 0.86 ± 0.01 | 0.87 ± 0.03 | 0.188 | No significative |
| Density index<br>(WE/SL) | All | 4.44 ± 1.32 | 4.48 ± 1.59 | 0.700 | No significative |
| | Portugal | 4.41 ± 0.61 | 4.43 ± 0.54 | 0.991 | No significative |
| | France | 4.47 ± 1.87 | 4.52 ± 2.03 | 0.784 | No significative |

**Figure 3.**
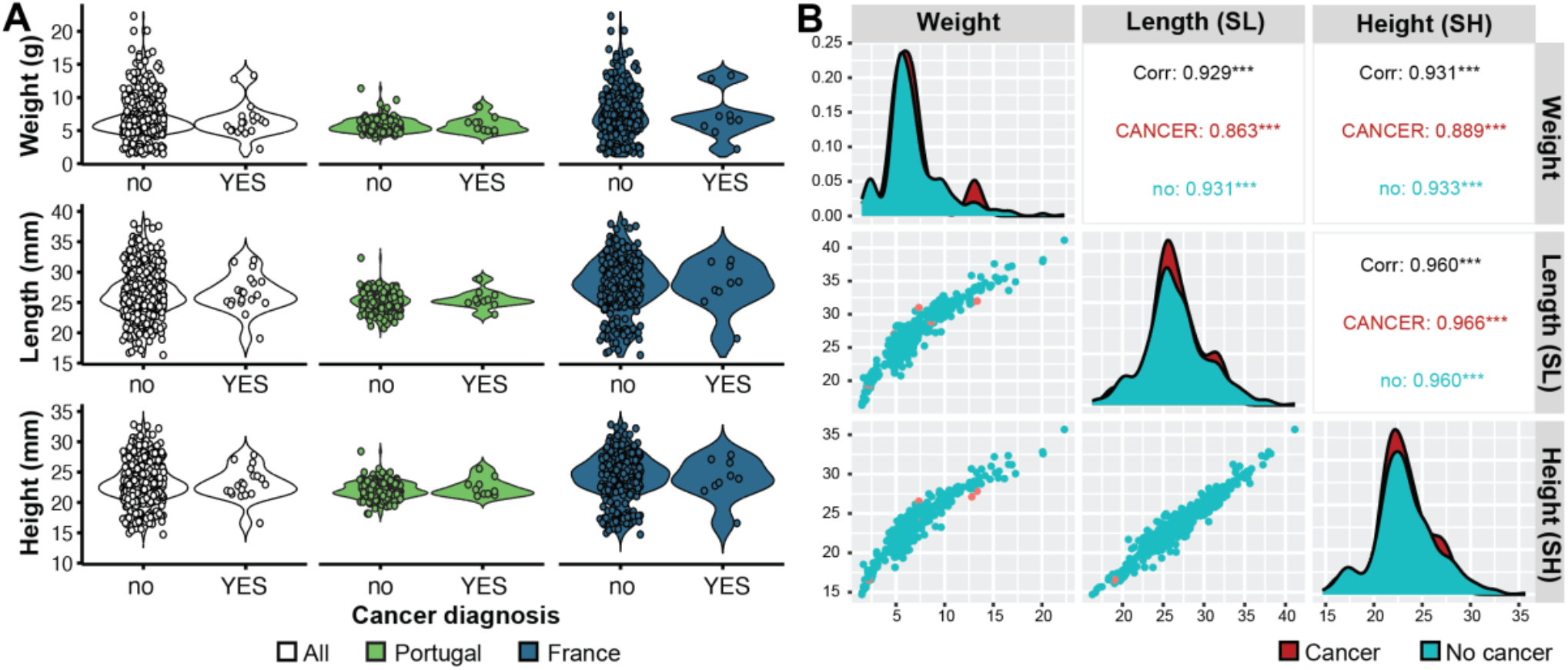
Morphometric analysis of cockles with and without cancer. **(A)** Violin plots showing the distribution of Weight, Length, and Height for all samples and separated by country (Portugal and France), comparing individuals diagnosed with cancer (yes) and without cancer (no). **(B)** Pair plot showing the relationships between Weight, Length, and Height coloured by diagnosis. The diagonal shows histograms for each variable, the lower panels display scatter plots, and the upper panels show Pearson correlation coefficients (Corr), both overall (black) and for each group. All variables are strongly correlated, though weight shows slightly lower correlations in the cancer group.

Correlation analyses revealed strong positive relationships between weight, length, and height in all individuals (Figure 3B). Although weight showed slightly lower correlations in the cancer group, suggesting greater variability in the relationship between weight and height, these differences were not statistically significant. That indicates that cancer did not substantially alter the proportional growth of these traits.

## 4. DISCUSSION

### 4.1. Spatiotemporal patterns of BTN distribution in cockles

BTN prevalence varies significantly across regions, species, and time, highlighting the complex interplay between host biology and environmental factors. In our study, we observed differences in BTN prevalence among locations, with the Algarve region presenting the highest infection rates (Table 1, Figure 2C). As all samplings were conducted in spring or summer, seasonal effects were minimized, but longer-term monitoring is needed to confirm temporal trends. Notably, the disease was found exclusively in cockles within the two studied areas, while other species sampled showed no signs of infection. Given the reports of cross-species transmission (Metzger *et al*., 2016; Yonemitsu *et al*., 2019; Hammel *et al*., 2021; Garcia-Souto *et al*., 2022), these results suggest that host-specific factors contribute to the observed patterns of disease distribution and the presence of cancer in a particular species in an area does not implicate cross-transmission to co-habiting species.

In the Algarve, both Ria Formosa and Fuzeta showed similar rates, whereas Alvor showed no signs of cancer in the sampled cockles. These findings might be due to the ecological distinctions between the two regions. Ria Formosa is considered a restricted or leaky coastal lagoon, whereas the Alvor system is considered a chocked system with only one narrow channel connecting the lagoon to the ocean which implies that water residence times in Ria Formosa are generally higher than in the Ria de Alvor (Brito *et al*., 2012). Given that cancer transmission requires cell transfer through water (Burioli *et al*., 2021; Giersch *et al*., 2022; Sunila and Farley, 1989), a minimum water residence time may be necessary for successful transmission.

In Normandy, the 6% prevalence in the Estuaire de l’Orne represents a significant rate contrasting with the neighbouring Brittany cases reported in 2017 where prevalence was 2.1% and 1.4% at Roscoff (Bruzos *et al*., 2023). Furthermore, in 2024, no cases were detected in Brittany’s Baie de Saint-Brieuc (0%). To our knowledge, there were no previous data available for Normandy, making this the first report in this region. It suggests either a recent onset of transmission or an undetected presence of the disease in Normandy. Similar to the Algarve, ecological differences among the sampling points may also play a role. The Baie de Veys and Estuaire de l’Orne are more strongly influenced by anthropogenic activities and fluvial inputs, whereas the Baie de Saint-Brieuc, a designated natural reserve, experiences lower anthropogenic pressure (Baffreau *et al*., 2017; Dabouinneau et *al*., 2015). These findings emphasize the need for cancer surveillance across other potential hot spots in northern France.

Temporal variations in BTN prevalence have been reported in previous studies, with seasonal and interannual fluctuations likely driven by environmental factors such as temperature, salinity, and nutrient availability (Díaz *et al*., 2016). There has been little agreement, however, as to the nature of the seasonal cycle or its cause (Carballal *et al*., 2015). These variables may influence both the susceptibility of host populations and the transmission dynamics of the cancer. In this study we compared BTN prevalence in Ria Formosa with previous reports of spring/summer collections over a time period of 7 years (2017-2024) finding a decrease on the prevalence potentially linked to a decrease on cockle populations. However, a direct causal relationship between disease prevalence and population decline cannot be established. It is possible that reduced host density limits transmission rates, contributing to lower prevalence, but other environmental or ecological factors may independently affect both disease dynamics and population trends. Additionally, sampling variability and potential detection biases should be considered when interpreting these patterns. Further long-term monitoring is required to determine if this temporal trend in the studied population is consistent.

Understanding the drivers behind these spatial and temporal variations is key for predicting future disease outbreaks and implementing effective management strategies.

### 4.2. Morphological variability of cancer cockles

Our findings suggest that hemic neoplasia in cockles does not affect the external shell morphology or animal weight, which contrasts with some studies pointing that some parasitic diseases in bivalves impact growth and shell morphology (Zannella *et al*., 2017). Advanced hemic neoplasia has been associated with significant physiological deterioration, often manifesting as gonad castration and mantle recession (Barber, 2004; Carballal *et al*., 2015). For example, Leavitt *et al*. (1990) found that *Mya arenaria* with advanced cases of disseminated neoplasia had significantly lower tissue dry weight and condition index than clams without the disease. Further studies with larger sample sizes are needed to conclusively determine if disease progression leads to significant reduced growth rate or weight loss in cockles.

Traditionally, morphological abnormalities or condition indices have been used as preliminary indicators of disease in bivalve populations (Zannella *et al*., 2017). However, our results suggest that external morphology cannot be a reliable diagnostic tool for detecting hemic neoplasia and its impacts on cockle populations. Instead, cytological examination, flow cytometry, or molecular diagnostics, are necessary for accurate disease surveillance (Giersch *et al*., 2022; Santamarina *et al*., 2025; Skazina *et al*., 2021).

### 4.3. Bivalve susceptibility to cancer transmission

Early observations of cancer appearance in bivalves after co-habitation (Appeldoorn, Brown and Chang, 1984), the successful transplantation of cancer cells (Elston, Kent and Drum, 1988), cancer cells survival in the seawater (Giersch *et al*., 2022; Sunila and Farley, 1989), and reports of interspecific cancer contagion, suggested that an area with a high prevalence of BTN in cockles such as the Algarve could be a location to originate interspecific contagions. However, after the inspection of 437 clams of 2 species, no cancer was found.

The absence of hemic neoplasia in *Ruditapes* spp. and *S. plana* in areas where BTN is present in cockles suggest that it may be restricted to cockles and raises questions regarding host specificity and resistance mechanisms. While some bivalves appear highly susceptible to BTN, others remain unaffected despite sharing the same habitat suggesting that environmental exposure alone is not sufficient for transmission (Díaz, Villalba and Carballal, 2017). Genetic predisposition and immunological differences between species likely influence their ability to resist or tolerate BTN infection, a hypothesis that requires further investigation.

This finding is reassuring for the Portuguese aquaculture industry, where about 90% of *Ruditapes* spp. are harvested in Ria Formosa (Silva *et al*., 2021). Thus, no immediate threat of cross-species transmission to these bivalves has been detected, although continued monitoring remains essential to identify any changes in disease patterns. However, given the limited sample size for this species in Normandy, our capacity to detect low-prevalence infections was constrained, and we caution against interpreting the absence of detected cases as evidence of true disease absence.

## 5. CONCLUSION

This study provides new insights into the prevalence and distribution of hemic neoplasia in cockles along the western coast of France and the southern coast of Portugal. The detection of the disease in Normandy expands its known range in France, while the observed spatio-temporal patterns in prevalence in the Algarve suggest potential environmental or population-related factors influencing disease dynamics. Despite previous reports of cross-species transmission in bivalves, no cases of hemic neoplasia were found in cohabiting *Ruditapes spp*. Or *S. plana*, suggesting species-specific susceptibility. Additionally, no significant differences in shell morphology or weight were observed between affected and non-affected cockles, indicating that external traits are not reliable indicators of disease presence. These findings highlight the need for continued monitoring to assess long-term trends in disease prevalence and better understand the factors influencing the spread and impact of hemic neoplasia in bivalve populations.

## Acknowledgments

We would like to thank Sr. Elio, Sr. Isidoro, Christophe Roger and Laurent Dabouinneau for providing technical assistance during field sampling. We also extend our thanks to Sushee Dunn (AquaExcel), Claire Dupree, Emilie Duval, Sandrine Soro, and Sylvia Forte (Université de Caen Normandie) for their administrative support. The images presented in Figure 1A-C were taken during sampling and processing activities conducted as part of this study. Additionally, we acknowledge the staff of CREC and CCMAR marine stations for their logistical support, and access to essential resources, which greatly facilitated our research.

This study was supported by the European Union’s Horizon 2020 research and innovation programme through the Marie Skłodowska-Curie grant agreement no. 101034329 and the Région de Normandie as part of the WinningNormandy fellowship awarded to ALB. The 2020 Algarve sampling data was collected under the European Research Council Starting Grant ‘Scuba Cancers’ (716290). This study received support from the Portuguese node of EMBRC-ERIC, specifically EMBRC.PT ALG-01-0145-FEDER-022121 and funding from the European Union’s Horizon 2020 research and innovation programme through project Aquaexcel 3.0 under grant agreement No 871108. This research also received support from the Exploration Fund Grant of The Explorers Club (2024 call).

## Supplementary material

- Bruzos_et_al_Supplementary Data 1.xlsx: Sampling information
- Bruzos_et_al_Supplementary Data 2.xlsx: Morphometrics data
- Bruzos_et_al_Supplementary Data 3.pdf: Cytological preparations used for diagnosis.
- Graphical Abstract

## Additional Information

### Data Accessibility

The datasets supporting this article have been uploaded as part of the Supplementary Material.

### Authors’ Contributions

A.L.B.: conceptualization, fieldwork, laboratory sample processing, data curation, formal analysis, visualization, funding acquisition, project administration, supervision, writing - original draft, writing - review and editing; A.M.: fieldwork, laboratory sample processing, data curation, formal analysis, visualization, writing - review and editing; G.B.: fieldwork, data curation, formal analysis, writing - review and editing; A.S.: fieldwork, writing - review and editing; A.V.F.: fieldwork, laboratory sample processing, data curation, writing - review and editing; A.M.A.: fieldwork, laboratory sample processing, funding acquisition, project administration, supervision, writing - review and editing; C.D.: funding acquisition, project administration, supervision, writing - review and editing. All authors gave final approval for publication and agreed to be held accountable for the work performed therein.

